# The volitional control of individual motor units is constrained within low-dimensional neural manifolds by common inputs

**DOI:** 10.1101/2024.01.05.573816

**Authors:** Julien Rossato, Simon Avrillon, Kylie Tucker, Dario Farina, François Hug

## Abstract

The implementation of low-dimensional movement control by the central nervous system has been debated for decades. In this study, we investigated the dimensionality of the control signals received by spinal motor neurons when controlling either the ankle or knee joint torque. We first identified the low-dimensional latent factors underlying motor unit activity during torque-matched isometric contractions. Subsequently, we evaluated the extent to which motor units could be independently controlled. To this aim, we used an online control paradigm in which participants received the corresponding motor unit firing rates as visual feedback. We identified two main latent factors, regardless of the muscle group (vastus lateralis-medialis and gastrocnemius lateralis-medialis). The motor units of the gastrocnemius lateralis could be controlled largely independently from those of the gastrocnemius medialis during ankle plantarflexion. This dissociation of motor unit activity imposed similar behavior to the motor units that were not displayed in the feedback. Conversely, it was not possible to dissociate the activity of the motor units between the vastus lateralis and medialis muscles during the knee extension tasks. These results demonstrate that the number of latent factors estimated from linear dimensionality reduction algorithms does not necessarily reflect the dimensionality of volitional control of motor units. Overall, individual motor units were never controlled independently of all others but rather belonged to synergistic groups. Together, these findings provide evidence for a low-dimensional control of motor units constrained by common inputs, with notable differences between muscle groups.

**Significance statement:** In this study, we initially examined the latent factors underlying motor unit activity in the vastii or gastrocnemii muscles during torque-matched isometric contractions. We then explored the extent to which these motor units could be controlled independently, using an online control paradigm where participants received visual information on motor unit firing rates. Although participants were able to dissociate the activity of a few motor unit pairs (from the gastrocnemius medialis and lateralis muscles), our results provide direct evidence of a low-dimensional control, constrained by common inputs, limiting flexibility in motor unit recruitment. Furthermore, we show that the number of latent factors identified by dimensionality reduction algorithms does not necessarily reflect the dimensionality of volitional control of motor units.

## Introduction

To generate movement, the central nervous system must provide appropriate inputs to the spinal motor neurons. The relatively large number of spinal motor neurons theoretically implies a control space of high dimensionality. How the central nervous system navigates through this high-dimensional space to generate neural commands remains an open question (Latash, 2021).

Observations from the motor areas of the cortex (Gallego et al., 2017) and the spinal cord (De Luca and Erim, 1994; Farina and Negro, 2015) have led to the view that the central nervous system relies on neuronal population coding rather than single neuron coding to control movements. This is supported by studies reporting correlations between the firing activity of spinal motor neurons that innervate the same or different muscles (Gibbs et al., 1995; De Luca and Erim, 2002; Keen et al., 2012). As the outputs of two motor neurons cannot be correlated without a certain degree of correlation in their inputs (Rodriguez-Falces et al., 2017), this correlated activity underlies the common inputs received by the motor neurons.

The control of motor neurons by common inputs aligns with the synergy hypothesis (Bernstein, 1947), which proposes that a small number of inputs project to a greater number of controlled structures through specific neural circuits (Bizzi and Cheung, 2013; Takei et al., 2017). In this view, large groups of motor neurons (synergies) would share common inputs that constrain their firing behavior within a low-dimensional neural manifold, i.e., a low-dimensional subspace in a state space in which each axis represents the activity of one motor neuron. Despite being computationally efficient, this strategy would impose some “rigidity” (Marshall et al., 2022) in the control of motor neurons receiving the same inputs, as they would always be activated in the same order according to the size principle (Henneman, 1957). An alternative hypothesis to hardwired neural synergies suggests that the low-dimensional control of motor neurons originates from task constraints (Tresch and Jarc, 2009). These task constraints, along with other factors such as musculoskeletal geometry, reduce the set of feasible activation patterns (Kutch and Valero-Cuevas, 2012). From this perspective, the distribution of common inputs to motor neurons would be task-dependent, leading to an apparent flexible control.

The rigid control imposed by hard-wired synergies implies the presence of neural constraints in the volitional control of motor units, making it impossible to independently control two motor units that receive a large proportion of common inputs. The volitional control of motor units has been tested using a biofeedback paradigm, pioneered by Basmajian (1963). This approach has yielded conflicting results, with some studies showing that the activity of motor units from the same muscle can be dissociated, with reversed recruitment order (Harrison and Mortensen, 1962; Formento et al., 2021); and others showing that such a volitional dissociation cannot be achieved (Bracklein et al., 2022). These discrepancies may be due to different underlying distributions of common inputs to motor units, a factor not systematically assessed by these studies. Furthermore, previous studies reporting volitional dissociation of single motor units (Harrison and Mortensen, 1962; Formento et al., 2021) did not assess behavior at the population level. This prevents determining whether these units were controlled independently of all others, thereby hindering a definitive conclusion regarding the true level of flexibility at the population level.

In this study, we hypothesized that the central nervous system mainly adopts a rigid control of the motor units during a single-joint task. We first assessed the low-dimensional latent factors underlying motor unit activity during isometric torque-matched tasks. Subsequently, we evaluated the extent to which the same motor units could be independently controlled using an online control paradigm in which participants received the corresponding motor unit firing rates as visual feedback. Because of the freedom to explore different inputs to motor neurons using the online motor unit control paradigm, this approach provides evidence of the flexibility or rigidity of the volitional control of motor units.

## Methods

### 1. Participants and ethical approval

Thirteen physically active male volunteers (mean±standard deviation; age: 29.6±6.9 years, height: 179±5 cm, and body mass: 73.3±4.9 kg) participated in the triceps surae experiment, and eight physically active male volunteers (age: 30.4±6.5 years, height: 176±6 cm, and body mass: 75.6±4.7 kg) participated in the quadriceps experiment. Three of them participated in both experiments. Participants had no history of lower leg pain with limited function that required time off work or physical activity, or a consultation with a health practitioner in the previous 6 months. The procedures were approved by the local ethics committee (CERNI – Nantes Université, n°04022022), and were performed according to the Declaration of Helsinki, except for registration in a database. The participants were fully informed of any risks or discomfort associated with the procedures before providing written informed consent to participate. Notably, we specifically recruited males, as the pilot data revealed that too few (if any) GL or VM motor units could be identified with sufficient accuracy in females. This aligns with previous works showing that a greater number of motor units can be decomposed in males than in females (although the reasons for this discrepancy are yet to be well established) (Del Vecchio et al., 2020).

### 2. Experimental design

We tested our hypothesis on populations of motor units, selecting them based on different anticipated levels of common inputs, as indicated by previous work using coherence analysis: i) motor units from the same muscle (gastrocnemius medialis; GM-GM), which receive a large proportion of common inputs (Hug et al., 2021), ii) motor units from different muscles (vastus lateralis and medialis; VL-VM), which receive a large proportion of common inputs (Laine et al., 2015), and ii) motor units from different muscles (gastrocnemius lateralis and medialis; GL- GM), which receive a small proportion of common inputs (Hug et al., 2021).

For the triceps surae experiment, the participants sat on a dynamometer (Biodex System 3 Pro, Biodex Medical, USA) with their hip flexed at 80°, 0° being the neutral position, and their right leg fully extended (Figure 1). Their ankle angle was set to 10° of plantarflexion (0° being the foot perpendicular to the shank). As foot position may impact the distribution of neural drive between the GL and GM muscles (Hug et al., 2021; Crouzier et al., 2024), the foot was securely maintained in neutral position. For the quadriceps experiment, the participants sat on the dynamometer with their hips flexed at 80° and the knee of their right leg flexed at 80°, 0° being the full extension (Figure 1). Inextensible straps were tightened during both tasks to immobilize the torso, pelvis, and thighs on the test side. Notably, these joint configuration, muscles, and types of contractions have been previously shown to lead to a high yield of decomposed motor units (Avrillon et al., 2021; Levine et al., 2023). Each experiment involved a single experimental session lasting approximately 3 h.

**Figure 1.**
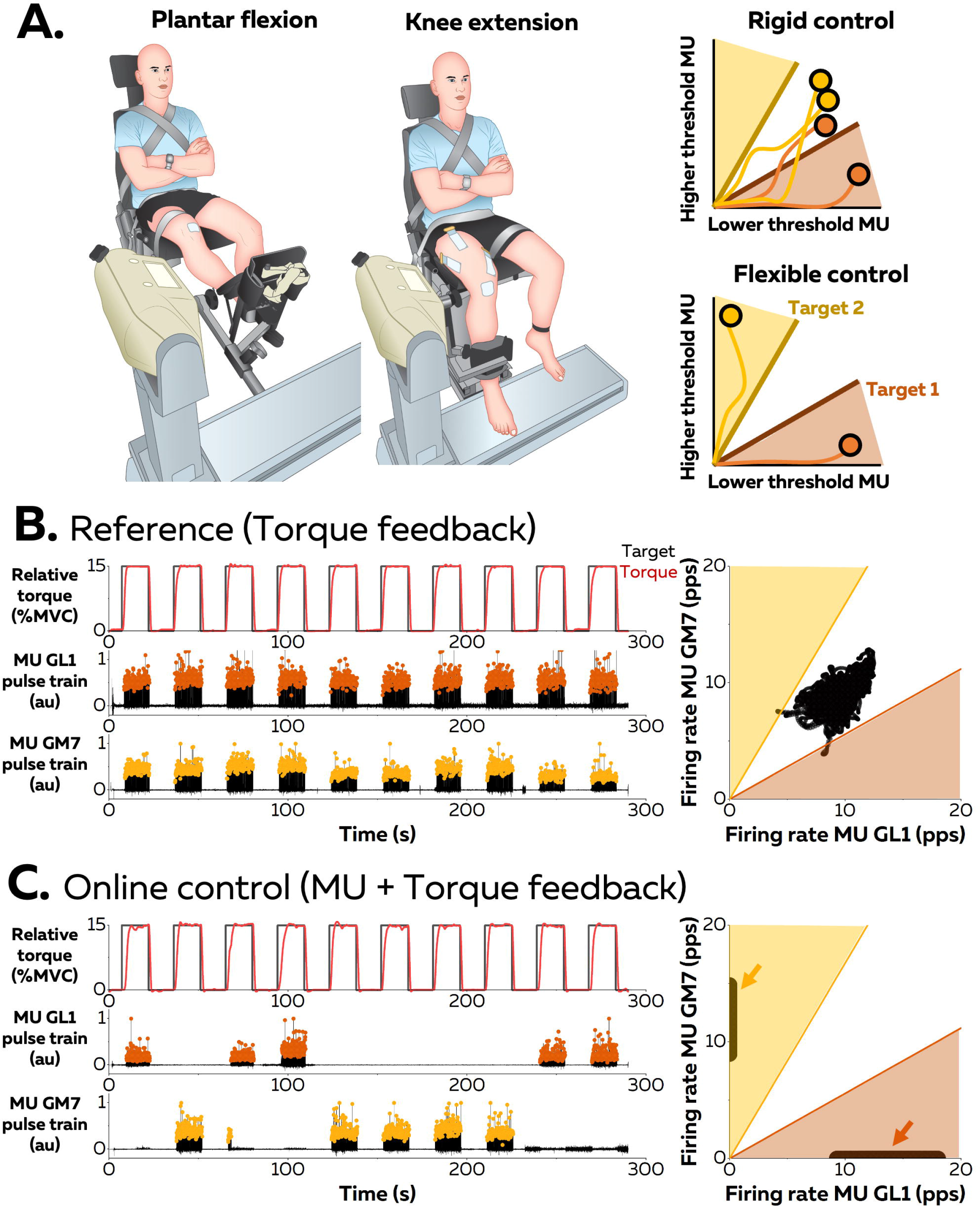
Experimental setup and protocol. (A) Participants performed submaximal isometric plantar flexions or knee extensions. Electromyographic signals were recorded using grids of surface electrodes and decomposed into motor unit discharge times. We used an online feedback control paradigm to let participants directly control motor units firing rates and test the hypothesis that this control is ‘rigid’ during a single-joint task. (B) Participants first performed a series of 10 contractions with torque feedback (reference contractions, B). In the second part of the experiment, motor unit firing rates were estimated in real-time and converted into visual feedback displayed to the participant (online control contractions, C). The visual feedback consisted of a two-dimensional space in which a cursor moved according to the firing rates of two motor units. Two targets (Target 1 and Target 2) were alternately displayed to the participant, who had to differentially modulate the firing rates of the two motor units to alternately move the cursor towards one of the two targets. (B) The firing rates of the motor units #1 from the Gastrocnemius lateralis (GL) and #7 from the Gastrocnemius medialis (GM) of one participant are displayed across the ten trials. Note that the two-dimensional space is displayed in Panel B but was not provided as feedback to the participants during the reference contractions. This example shows a clear dissociation of the activity of two motor units from different muscles, i.e. GL and GM, during the online control. Such a dissociation was not observed for GM-GM and Vastus lateralis-vastus medialis motor units.

The protocol was the same, regardless of the experiment (triceps surae or quadriceps). The experimental session began with a standardized warm up, which included five 3-s isometric contractions at 50%, 60%, 70%, and 80% and three 3 s contractions at 90% of the participants’ subjective maximal torque. After 2 min of rest, the participants performed three maximal voluntary contractions (MVC) for 3 s, with 60 s of rest in between. Peak MVC torque was considered as the maximal value obtained from a moving average window of 250 ms. Participants then performed one trapezoid isometric contraction (10 s ramp up, 30 s plateau, and 10 s ramp down) at a submaximal torque to identify motor unit filters offline (referred to as “baseline” contractions). For the triceps surae protocol, the torque level was chosen on an individual basis based on the number of identified motor units. Specifically, in some participants, the GL muscle was minimally activated at the lowest torque levels (15% and 20% MVC), leading to no or a few identified motor units. In this case, a higher torque level (20 or 25% MVC) was tested. The retained torque was 15% MVC for five participants, 20% MVC for four participants, 25% MVC for one participant. This torque level was then kept the same during the whole experimental session. For the quadriceps protocol, the torque level was set at 10% MVC as it was sufficient to recruit motor units in both muscles. The choice of a relatively low torque level was also motivated by the need to limit fatigue, which develops faster during knee extension than plantarflexion (Rossato et al., 2022). After a 15-30 min rest period during which automatic decomposition and manual editing of the motor units were performed, the participants performed ten submaximal isometric contractions for 15 s, with 15 s of rest in between. Only torque feedback was provided to the participants during these contractions (referred to as “reference” contractions) so that they could be used to assess the “natural” motor unit recruitment strategies.

The rest of the experimental session consisted of two to four blocks of a series of submaximal isometric contractions during which the participants were instructed to dissociate the activity of two motor units while keeping the plantarflexion torque constant. Specifically, participants navigated a cursor in a 2D space were the *x-* and *y-* axes represented the firing rate of motor units #1 and #2. The scale of each axis was set up from 0 to a maximum of two times the mean firing rate of the corresponding motor unit measured during the baseline contraction. This ensured that the cursor was not biased toward any of the axes during natural contractions. A feedback of torque output was also provided to the participants in the form of a vertical gauge that the participants had to keep between two lines representing ±5% of the target torque. For the triceps surae protocol, we aimed to provide visual feedback of four different pairs of motor units for each participant: two pairs composed of motor units of both the GM and GL (between-muscle pairs) and two pairs composed of motor units of the GM (within-muscle pairs). For the quadriceps experiment, we aimed to provide visual feedback on two different pairs of motor units composed of the motor units of both the VM and VL (between-muscle pairs). Each pair was tested separately within a block of contractions as described below. Of note, because of either a decrease in the accuracy of motor unit identification during the experiment or the overall long duration that made the lower limb position progressively uncomfortable, we tested fewer pairs of motor units (mean number of tested pairs for each participant: 3.3 ± 0.9 pairs [total = 17 for GM-GL and 16 for GM-GM] for the triceps surae experiment and 1.8 ± 0.4 pairs [total = 9] for the quadriceps experiment). Notably, because a drastic drop in the decomposition accuracy across the experiment in some participants, these pairs originated from ten and five participants for the triceps surae and quadriceps experiment, respectively.

Each block started with an exploration phase during which the participants were instructed to navigate the cursor inside the 2D space during three 60 s periods, with 60 s of rest in between. During this exploration phase, participants were asked to explore the entire 2D space; that is, to alternately move the cursor close towards (or on) the *x* and *y* axis while maintaining the torque at the predetermined torque target. Notably, to move the cursor close to an axis (either *x* or *y*), the participants had to differentially change the firing rate of the two motor units (i.e., an increase in one unit with no change or a decrease in the other unit). For the subsequent contractions, the target spaces were added to the 2D space. These targets were in the shape of a triangle with one corner on the coordinate origin and two sides being one axis (*x* or *y*), and a line forming an angle of 30° with this axis (Figure 1). First, participants performed a series of ten 15 s contractions during which they were asked to move the cursor in the target space close to one of the axes (*x* or *y* [randomized]). The participants then performed another series of ten 15 s contractions during which they were asked to move the cursor in the target space close to the other axis. Finally, they performed a last series of ten 15 s contractions during which they were asked to move the cursor in the target space close to either the *x* axis (five contractions) or the *y* axis (five contractions). During this last series, the order of the target presentation (i.e., *x* or *y*) was randomized and unknown to the participants. The contractions were interspaced by a 14-s recovery period and the three series were interspaced by a 2 min rest period. The participants were informed that a successful trial was one in which the cursor stayed in the target space while producing the required torque, and that the ultimate objective was to move the cursor on the x or y axis, meaning that they kept one motor unit active while keeping the other motor unit off. Of note, for the purpose of this study only the third series of 10 contractions was analyzed (referred to as “online” contractions), the previous series being considered as training.

### 3. High-density surface electromyographic recordings

For the triceps surae experiment, high-density surface electromyographic (EMG) signals were recorded from four lower limb muscles: GM, GL, Tibialis anterior (TA), and VL. The TA and VL muscles were recorded to explore possible strategies for modulating the activity of the gastrocnemius motor units, such as reciprocal inhibition. Indeed, the TA and VL act as antagonists of the gastrocnemius in two main functions, i.e., plantarflexion for TA and knee flexion for VL. This was further motivated by a previous study that showed that adding knee extensor activity to plantar flexion led to a differential change in activation of the GM and GL muscles (Suzuki et al., 2014). For the quadriceps experiment, signals were recorded from two lower limb muscles: the VM and VL.

A two-dimensional adhesive grid of 64 electrodes (13×5 electrodes with one electrode absent in a corner, gold-coated, inter-electrode distance: 8 mm; [GR08MM1305, OT Bioelettronica, Italy]) was placed over each muscle. Before electrode application, the skin was shaved and cleaned with an abrasive gel (Nuprep, Weaver and company, USA). The adhesive grids were held on the skin using bi-adhesive foam layers (OT Bioelettronica, Italy). Skin-electrode contact was ensured by filling the cavities of the adhesive layers with a conductive paste. A 10- cm-wide elastic band was placed over the electrodes with a slight tension to ensure that all the electrodes remained in contact with the skin throughout the experiment. A strap electrode dampened with water was placed around the contralateral ankle (ground electrode) and a reference electrode (5 × 5 cm, Kendall Medi-Trace™, Canada) was positioned over the tibia (triceps surae experiment) or the patella (quadriceps experiment) of the ipsilateral limb. The EMG signals were recorded in monopolar mode, bandpass filtered (10-500 Hz) and digitized at a sampling rate of 2048 Hz using a multichannel acquisition system (EMG-Quattrocento; 400-channel EMG amplifier, OT Biolelettronica, Italy).

We used an open-source software developed and validated by our team to decompose the EMG signals in real-time (Rossato et al., 2023). First, EMG signals were recorded during a submaximal contraction, such that channels with artifacts and low signal-to-noise ratio could be removed from all further analyses. Second, the EMG signals collected during the baseline contraction were decomposed offline using a blind-source separation algorithm (Negro et al., 2016). This baseline contraction aimed to identify the separation vectors (motor unit filters). As highlighted in our previous validation work (Rossato et al., 2023), the accuracy of the online decomposition can be improved by manually editing the offline decomposition results. Therefore, we performed a manual editing step that consisted of i) removing the spikes close to the noise level, ii) recalculating the motor-unit filter and reapplying it over the signal, iii) adding new spikes recognized as motor unit firings, and iv) recalculating the motor-unit filter (Hug et al., 2021). For the contractions requiring real-time decomposition, EMG signals were transmitted by packages of 256 data points (125 ms with a sampling frequency of 2048 Hz), and the motor unit filters were applied over the incoming segments of the EMG signals. In addition of the incompressible delay of 125 ms, the computational time required by the software to display visual feedback is typically less than 15 ms (Rossato et al., 2023). We calculated the firing rate (i.e., spikes per second) of individual motor units as the sum of the spikes over a moving window of eight consecutive epochs of 125 ms). While this approach introduced a delay of 500 ms in the estimation of the firing rate, it allowed for smooth visual feedback, easier to control than an instantaneous estimate of firing rate.

### 4. Force measurements

For the triceps surae experiment, the dynamometer (Biodex System 3 Pro, Biodex Medical, USA) was equipped with a six-axis force sensor (Sensix, France). Even though the feedback provided during the experiment was based only on the effective plantarflexion torque, we considered the two other torque components (abduction/adduction and inversion/eversion) in the analysis to determine whether the dissociation of motor unit activity was associated with changes in mechanical output. These mechanical signals were digitized at a sampling rate of 2048 Hz using a multichannel acquisition system similar to that used for EMG (EMG-Quattrocento; 400-channel EMG amplifier, OT Biolelettronica, Italy).

### 5. Data analysis

#### Offline manual editing

To address the aims of this study, we analyzed 10 reference contractions (with torque feedback only) and 10 online contractions during which participants were instructed to move the cursor in the target space close to either the *x* axis (five contractions) or the *y* axis (five contractions). We identified 10.6 ± 3.8 (GM), 3.4 ± 1.5 (GL), 6.0 ± 1.9 (VL), and 2.2 ± 0.7 (VM) motor units per participant for the reference contractions and 9.8 ± 4.2 (GM), 2.1 ± 1.0 (GL), 4.8 ± 1.2 (VL), and 1.8 ± 0.8 (VM) motor units per participant for the online contractions. Notably, we not only analyzed the pairs of units used for the feedback, but also analyzed all the motor units that were identified during the baseline contraction. Each motor unit spike train was manually edited as described above.

#### Recruitment threshold

We estimated the recruitment threshold of each motor unit from the baseline contraction, which included a 10-s ramp up phase. The average of force over a 10-ms window centered around the first discharge time during the ramp phase was considered as an estimate of the motor unit recruitment threshold.

#### Dimensionality reduction

We used non-negative matrix factorization (Lee and Seung, 1999) to determine the low-dimensional latent factors underlying the behavior of motor units during the 10 concatenated reference contractions (i.e. torque-matched isometric contractions). The binary vectors, with discharge times equal to one, were convolved with a 400-ms Hanning window and normalized between 0 and 1. The smoothed discharge rates were then concatenated in a *p*-by-*n* matrix, where *p* is the number of motor units and *n* is the number of time samples. To select the number of latent factors, we fitted portions of the curve of the variance accounted for (VAF) with straight lines by iteratively decreasing the number of points considered in the analysis. The number of latent factors was considered as the first data point where the mean squared error of the fit falls under 5e-4 (Cheung et al., 2005; d’Avella et al., 2011).

As we selected two latent factors for both the triceps surae and the vastii muscles (see results), each motor unit was positioned in a two-dimensional space, where x and y coordinates were the weights of each motor unit in the two latent factors. As proposed by Levine et al. (2023), we estimated the level of common inputs for each pair of motor units by calculating the angle between their vectors (Figure 2). We considered that the higher the angle between two vectors, the lower the proportion of inputs shared between these units.

**Figure 2.**
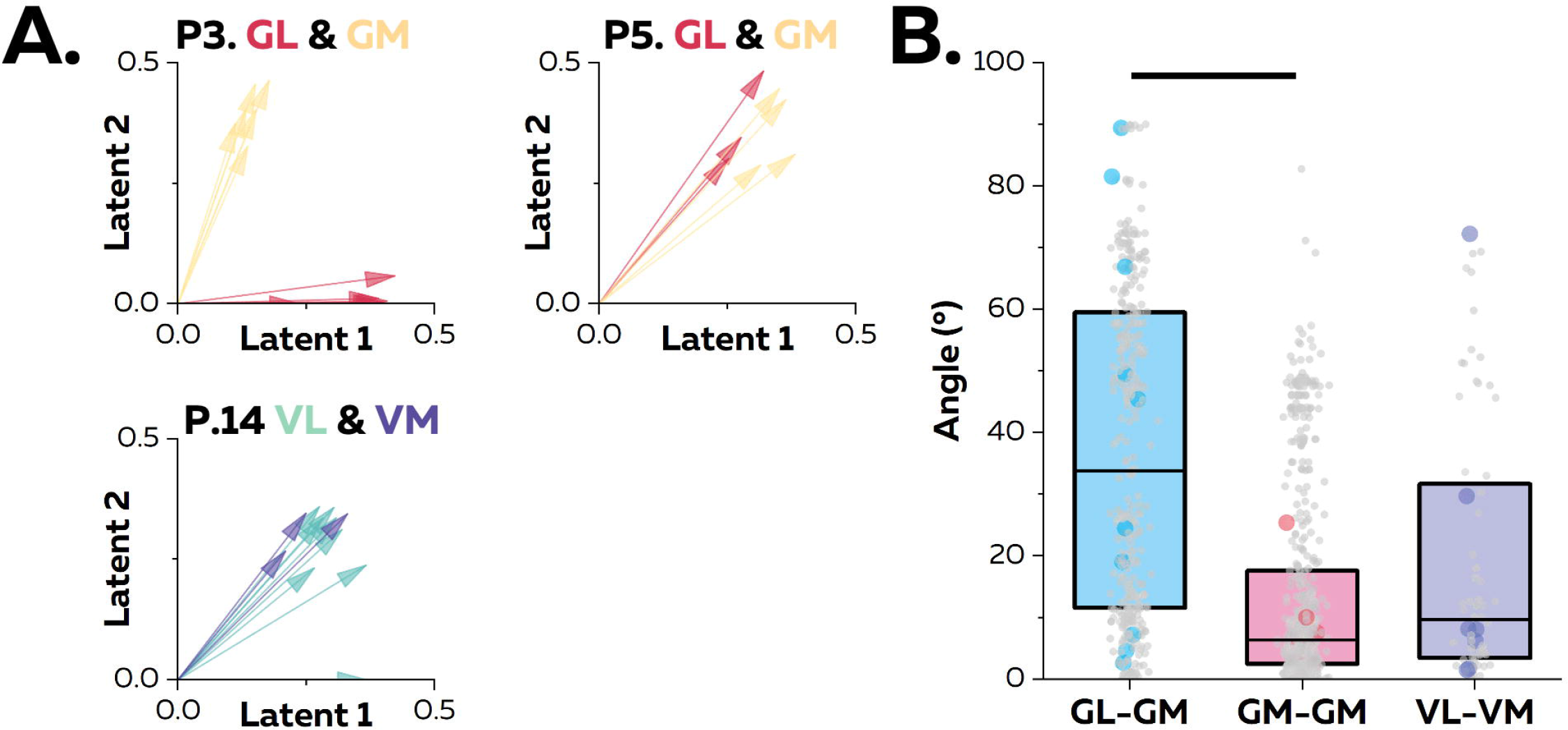
Motor unit behavior during the reference contractions. (A) Biplots in which the coordinates of each motor unit represent its weights within the two latent factors, as identified using non-negative matrix factorization (data depicted for three participants: P3, P5, and P14). Angles between vectors were calculated to estimate the level of common inputs for each pair of motor unit. It is worth noting that the firing activity of GL-GM motor units was either well correlated (P5) or uncorrelated (P3), while the firing activity of VL-VM motor units systematically covaried (P14). (B). Results are displayed for all participants, where each grey dot represents a pair of motor units and each colored dot represents a pair that was used as feedback in the second part of the experiment. The box denotes the 25^th^ and 75^th^ percentiles of the values distribution. The line is the median. The thick horizontal black lines denote a significant statistical difference between muscles (p<0.05).

#### Volitional dissociation of motor unit activity

For analysis purposes, the firing rate of the motor unit with the lowest recruitment threshold was depicted in the *x*-coordinate (Target 1) and the firing rate of the motor unit with the highest recruitment threshold was depicted in the *y*-coordinate (Target 2). To determine whether the participants were able to dissociate the activity of motor units, we calculated the success rate as the percentage of time spent within the required target area out of the total duration of the contraction. Notably, only data obtained while the participants reached the required torque level +/- 5% were considered in this calculation.

Three extreme scenarios were possible (Figure 1): i) the participant failed to reach Targets 1 and 2 (success rates: 0% and 0%), ii) the participant only reached Target 1 but failed to reach Target 2 (total success rates: 100% and 0%), and iii) the participant reached Targets 1 and 2 (success rate: 100% and 100%). While the second scenario may appear as a flexible control, it corresponds to the selective recruitment of the lowest threshold motor unit (cursor in Target 1), and therefore does not provide evidence of the ability to reverse the recruitment order. In the third scenario, participants succeeded in volitionally dissociating the activity of the two motor units. Specifically, this implies that they isolated the activation of the highest threshold unit, an observation that is inconsistent with a rigid strategy that would associate common inputs with the size-recruitment principle. Therefore, we considered that the first two scenarios supported the presence of common synaptic inputs constraining the firing rate of the two motor units, whereas the third scenario supported the presence of different inputs to the motor units.

#### Correlation between smoothed motor unit firings

To determine whether the behavior of the motor units displayed in the feedback imposed similar behavior on the non-displayed units, we calculated the correlation between the activity of each motor unit provided with feedback and that of each non-displayed unit. Specifically, we calculated the correlation coefficient between the smoothed firing rates over the 10 concatenated contractions of the online contractions using a cross-correlation function (max lag = 100 ms). The discharge times of each motor unit were first converted into continuous binary signals with ‘ones’ corresponding to the firing instances. The smoothed firing rate was obtained by convolving the binary vectors with a Hanning window of 400 ms.

#### EMG amplitude

To estimate the activation level of the TA and VL muscles during the triceps surae experiment, the 64 EMG signals recorded in a monopolar configuration were differentiated along the column direction to obtain 59 single-differential signals. The average rectified values (ARV) of each signal were estimated over a sliding window of 250 ms. The ARV of the 59 channels were finally averaged at each timestamp. ARV were averaged while each participant reached that target favoring the recruitment of motor units from the GL, from GM, or in between.

### 6. Statistical analysis

All statistical analyses were performed with RStudio (USA). First, quantile-quantile plots and histograms were used to check the normality of the data distribution. All the distributions were found normal except one: angle between vectors presented on Figure 2. In that case, data were log transformed. All the analyses were then performed using linear mixed effect models, implemented in the R package *lmerTest,* with Muscle (GL-GM, GM-GM, VL-VM) as a fixed factor and Participants as a random factor. When necessary, multiple comparisons were performed using the R package *emmeans*, which adjusts the *p*-value using the Tukey method. The significance level was set at p<0.05. Values are reported as mean ± standard deviation.

### 7. Code and Software accessibility

We used open-source software developed by our team to decode the activity of motor units in real-time (https://github.com/simonavrillon/I-Spin) and to edit the results (https://github.com/simonavrillon/MUedit). In addition, the entire dataset (raw and processed data) is available online (https://figshare.com/s/68b5a9b6dfd8e412b01c).

## Results

### Behavior of motor units during contractions with torque feedback

During the reference contractions, participants accurately tracked the target torque with a root mean square error of 0.4 ± 0.1% (plantarflexion) and 0.3 ± 0.1% (knee extension). We identified a total number of 40 (GL), 110 (GM), 34 (VL), and 15 (VM) unique motor units.

We applied non-negative matrix factorization to identify the main latent factors underlying motor units’ activity. When considering the GL and GM muscles, the analysis revealed one latent factor in two participants, two latent factors in six participants, and three latent factors in two participants. For the VL and VM muscles, two latent factors were identified in four participants, with one factor found in the remaining participant. An average of 2.0 and 1.8 latent factors was thus identified for GL-GM and VL-VM, respectively. Consequently, we decided to retain two latent factors for all participants for subsequent analysis, which accounted for a VAF of 82.2 ± 5.9 for GL-GM and 82.4 ± 3.1% for VL-VM.

For each participant, we displayed the motor units in a two-dimensional space, where x and y coordinates corresponded to the weight of each motor unit within each latent factor (Figure 2). The angle between two vectors, that is, two motor units, was calculated to estimate their level of common inputs. There was a main effect of Muscle (F(2, 40) = 113.2; p<0.001), with GM- GM (13.8 ± 16.5°) exhibiting lower angles than GL-GM (36.7 ± 26.0°; p<0.001) motor unit pairs. There was no difference between GM-GM and VL-VM (19.4 ± 21.7°, p=0.17) or between GL-GM and VL-VM motor unit pairs (p=0.34). To determine whether these results reflect neural constraints on motor unit firing activity, participants were asked to volitionally dissociate the firing rates of motor units in the second part of the experiment.

### Online control of motor unit firing rate

The participants were instructed to perform submaximal 15-s online contractions at the same intensity as the reference contractions while receiving additional visual feedback on the firing rates of two motor units (Figure 3). Even though differences in recruitment threshold between muscles need to be interpreted with caution due to force sharing strategies, it is worth noting that the pairs of motor units displayed to the participant in the different conditions exhibited differences in recruitment thresholds: 6.7 ± 4.2 %MVC (n=17) for GL-GM, 3.1 ± 1.7 %MVC (n=16) for GM-GM, and 2.0 ± 1.7 %MVC (n=9) for VL-VM.

**Figure 3.**
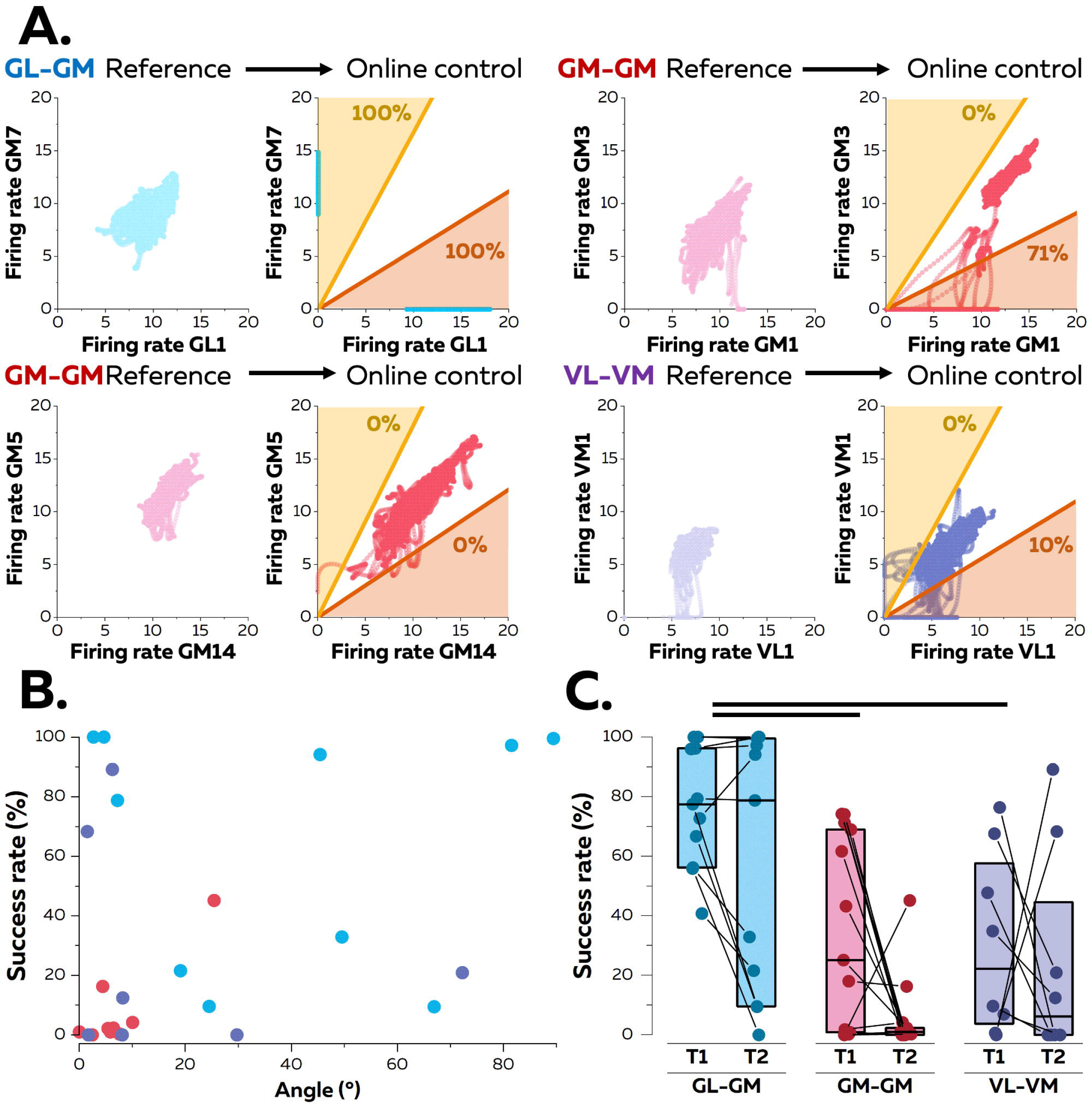
Volitional control of motor unit firing rates. (A) Typical examples of cursor positions are depicted for the 10 reference contractions (only torque feedback) and the online contractions (5 for each target). The success rate is displayed for each target. It is calculated as the percentage of time spent within the target (while maintaining the torque constant) over the total duration of the trial. (B) Relation between the angle between motor units estimated from the reference contractions and the success rate in target 2 (true flexible control) for all the displayed motor units. Each data point is a pair of motor units, and the color code is the same as in E. (C) Success rate for Targets 1 (T1) and 2 (T2). Each data point is a pair of motor units, the box denotes the 25^th^ and 75^th^ percentiles of the distribution of scores, and the line is the median. The thick horizontal black lines denote a significant statistical difference between muscles (p<0.05).

When considering the success rate, there was a main effect of Muscle (F(2, 28) = 16.3; p<0.001) and Target (F(1,47) = 6.0; p=0.018) but no interaction between Muscle and Target (F(2,47) = 0.8; p=0.48). Specifically, the success rate was higher for GL-GM than for both GM-GM (p<0.001) and VL-VM (p=0.008) regardless of the target. In addition, the success rate of Target 1 (lowest threshold unit) was higher than that of Target 2 (highest threshold unit) (p=0.018), regardless of the Muscle (Figure 3). Of note, four GL-GM motor unit pairs (recorded from two participants) exhibited a success rate close to 100% for both targets, demonstrating the ability of these two participants to selectively activate these units (Figure 3). In contrast, when considering GM-GM or VL-VM motor unit pairs, the success rate was close to zero for Target 2, which corresponded to a failure to selectively recruit the highest threshold unit. It is worth noting that three pairs (one for GM-GM and two for VL-VL) exhibited substantial success rates (>40%) for Target 2. However, this does not fully support a volitional independent control as the same pairs exhibited a success rate close to zero for Target 1. This can be explained by the fact that the difference in recruitment threshold between these units was very small (<2% MVC). Indeed, because of synaptic noise, the order of recruitment between units with similar recruitment thresholds may vary across contractions (Heckman and Enoka, 2012). It is also worth noting that some participants managed to activate only the motor unit with the lowest recruitment threshold (Target 1) but were not able to selectively activate the motor unit with the highest recruitment threshold (Target 2) (Figure 3). As mentioned previously, this observation does not imply the presence of independent inputs to the motor units. Interestingly, to selectively recruit the motor unit with the lowest threshold while maintaining the effective joint torque, the participants compensated with other muscles. For example, when feedback was provided on two GM motor units, they increased GL activation to limit the recruitment to the lowest threshold GM motor unit. Together, these results demonstrated that participants were only able to volitionally dissociate the activity of motor units from the GM and GL, which supports the assumption that motor units from the GM and GL receive different inputs.

The angle between motor units estimated from the reference contractions was not significantly correlated with the success rate in target 2 for either GM-GL (r=0.01; p=0.96) or VL-VM (r=- 0.21; p=0.62). Although the correlation was significant for GM-GM (r=0.84, p=0.002), this was mainly explained by one pair (Figure 3).

### Behavior of motor units at the population level during the online contractions

We determined whether the dissociation of motor unit activities imposed similar behavior on the motor units that were not displayed in the feedback, and therefore, from which activity was not constrained. For each motor unit provided with feedback, we calculated the correlation between its activity and that of each non-displayed unit (Figure 4). We found significant effects of Condition (F(1, 463) = 17.1; p<0.001) and Muscle (F(2, 465) = 134.4; p<0.001) with a significant interaction between Condition and Muscle (F(2, 463) = 55.2; p<0.001). Specifically, the correlation coefficients were significantly lower during the online control contractions than during the reference contractions for GL-GM motor unit pairs (0.42±0.28 vs. 0.78±0.17; p<0.001), while they did not change for GL-GL (0.87±0.16 vs. 0.87±0.14; p = 1.00) or for GM- GM pairs (0.90±0.12 vs. 0.89±0.18; p = 1.00; Figure 3B). The same analysis was performed for the VL-VM motor unit pairs. We found significant effects of Condition (F(1, 134) = 22.6; p<0.001) and Muscle (F(2, 136) = 3.2; p=0.042), but no significant interaction between Muscle and Condition (F(2, 134) = 2.7; p=0.073). Specifically, the correlation coefficients were lower during the online control contractions than during the reference contractions for all the pairs of motor units (0.89±0.14 vs. 0.98±0.02; p<0.001). However, it is worth noting that the correlations stayed at a high level, regardless of the condition.

**Figure 4.**
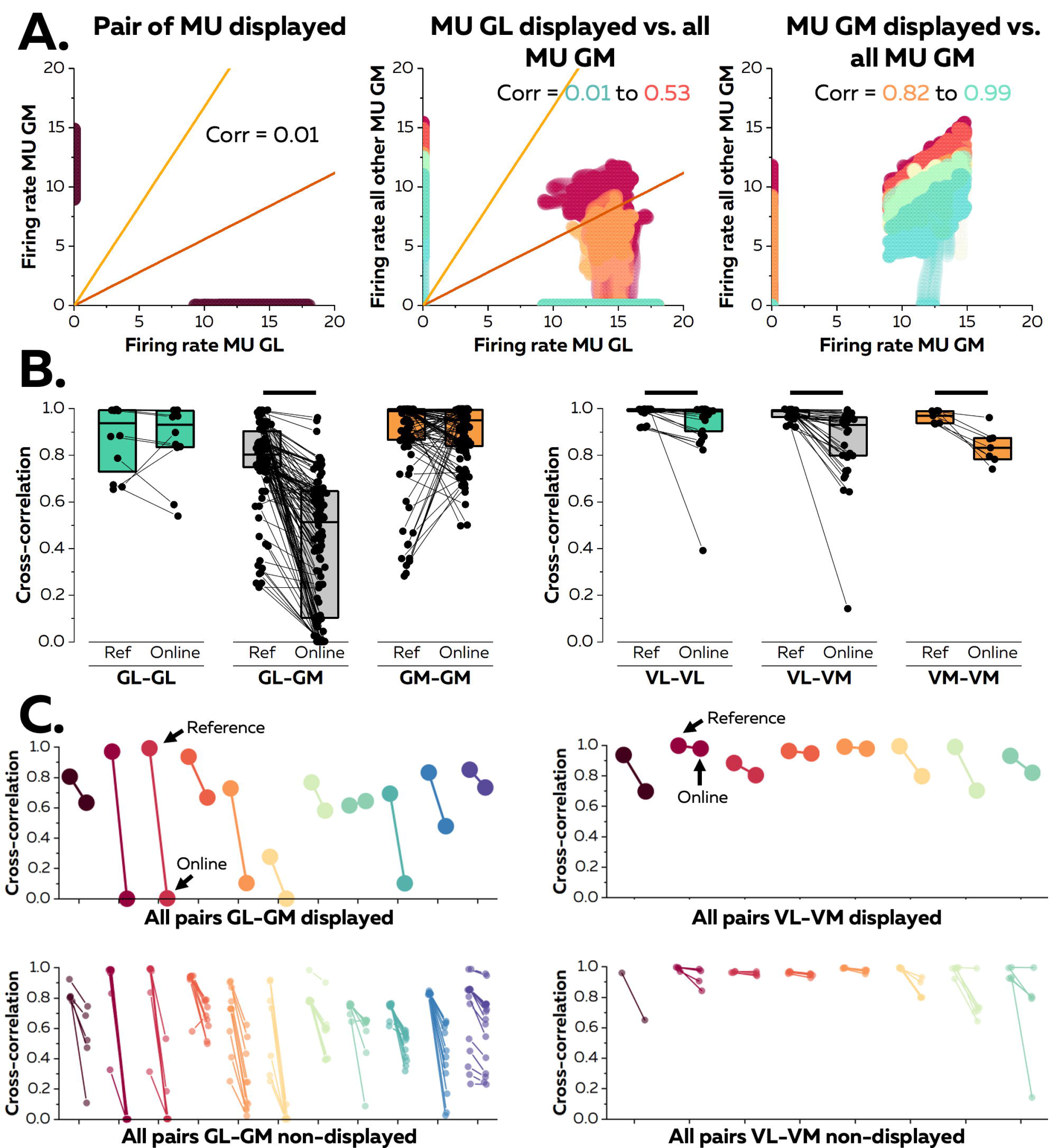
Changes in firing rate at the motor unit population level. (A) We calculated the correlation coefficients between the smoothed firing rates of the motor units displayed to the participant and the smoothed firing rates of the non-displayed units. In this example, the correlation coefficient between the firing rates of the two displayed motor units was 0.01 (left panel), showing a perfect dissociation of the activity of the two units. We extended the analysis to the correlation between the firing rate of the displayed motor units from the GL (central panel) or GM (right panel) and all the other motor units from GM. Each pair of motor units has a different color. (B) Correlation coefficients between the motor units (displayed and non-displayed to the participants) during the reference contractions and the online control contractions. Each data point is a pair of motor units, and the connecting line denotes the change in correlation between conditions. The box denotes the 25^th^ and 75^th^ percentiles of the distribution of coefficients and the black line is the median. The thick horizontal black lines represent a significant statistical difference between conditions. (C) The changes in correlations observed between reference contractions and online control trials were separated between all pairs of motor units displayed to the participant (top panel). The changes in correlations between the displayed units and those non-displayed units are shown in the bottom panel.

Overall, these results highlighted that uncoupling the firing rates of pairs of GL-GM motor units resulted in similar behaviors at the population level (Figure 4). Notably, 81.9 ± 40.2% of the non-displayed GL-GM pairs of motor units mirrored the trend observed in the displayed GL- GM pairs of motor units. By extension, non-displayed motor unit pairs from the same muscle (GL-GL or GM-GM; Figure 3B) remained highly correlated.

### Strategies associated with dissociation of motor unit activity during the online task

We examined the control strategies when the participants aimed to dissociate the activity of the GL-GM motor units. First, we calculated the ankle moments during the reference contractions and the online control contractions while the cursor was in the target favoring GL recruitment (TGL), GM recruitment (TGM), or the position in between (Tgas) (Figure 5). There was no main effect of Condition on ankle moments in the Dorsiflexion/Plantarflexion (F(3, 31) = 0.04; p=0.99) or Inversion/Eversion axes (F(3, 31) = 1.2; p=0.31). Conversely, a significant effect of Condition was found for the ankle moment in the Abduction/Adduction axis (F(3, 31) = 4.1; p=0.014), showing that selectively recruiting the GM motor unit was associated with more abduction (-10.9±13.0 N.m), and conversely selectively recruiting the GL motor unit was associated with more adduction (7.4±17.7 N.m; p=0.015, Figure 5). No other significant differences were observed (all p > 0.068).

**Figure 5.**
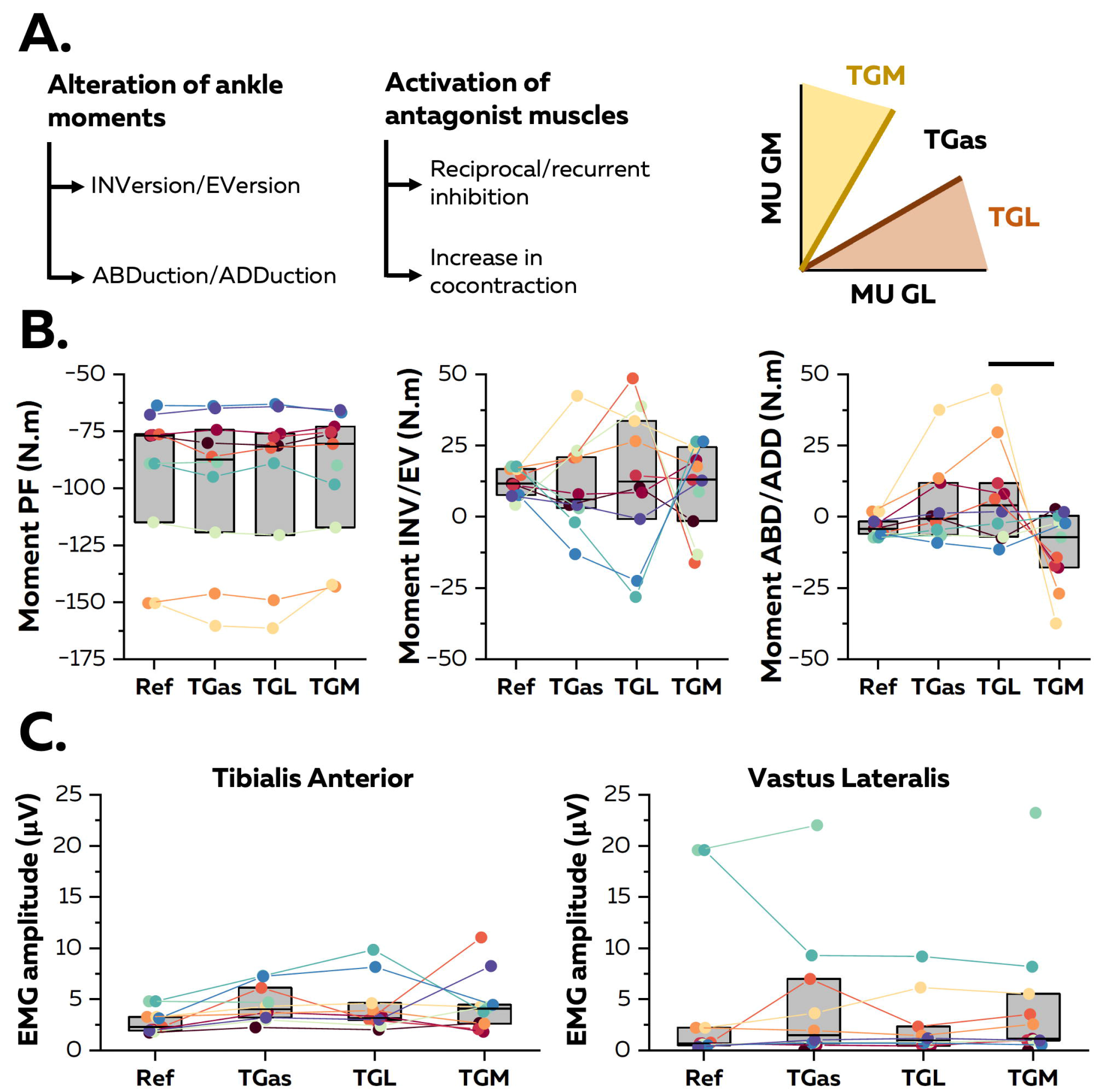
Control strategies during the online task. (A) To investigate the strategies associated with the dissociation of GL and GM motor unit firing rates, we identified the time stamps where the cursor was in the target favoring GL (TGL), or GM (TGM) motor unit recruitment. TGas refers to a cursor position between the two targets. Ankle moments and EMG amplitudes of the Tibialis anterior and Vastus lateralis were averaged over these periods and compared. (B) Average ankle force moments during the reference (Ref) contractions and the three targets. Each data point is a participant and the connect lines denote the variations between conditions. The box represents the 25^th^ and 75^th^ percentiles of the distribution of moments and the black line is the median. (C) Each data point is a participant and the connecting lines denote the variations between conditions. The box represents the 25^th^ and 75^th^ percentiles of the distribution of EMG amplitudes and the black line is the median. The thick horizontal black lines represent a significant statistical difference between conditions (p<0.05).

Second, we tested whether the participants increased the activation of their antagonist muscles (Tibialis Anterior [TA], Vastus Lateralis [VL]), which could have increased co-contraction around the ankle joint and modulated reciprocal/recurrent inhibition (Suzuki et al., 2014; Yavuz et al., 2018). This was specifically important for distinguishing between the central and peripheral origins of the observed dissociation of the GM and GL motor units. There was no main effect of Condition for TA (F(3, 31) = 2.1; p=0.12) or VL (F(3, 31) = 0.2; p=0.87; Figure 5), suggesting that dissociation of the GL and GM motor units was not achieved through modulating antagonist muscle activity.

## Discussion

We employed an online control paradigm in which participants were asked to volitionally dissociate the firing activity of pairs of motor units. In most cases, participants were able to fully dissociate the activity of pairs of GL-GM motor units, but this strategy imposed a similar behavior at the population level, that is, dissociation of the firing rate was observed for all other GL-GM motor unit pairs that were not displayed as feedback. Conversely, volitional dissociation of firing rates was not observed between motor units of other anatomically defined synergist muscles (VL and VM) or between motor units of the GM muscle. Together, our results provide evidence of a low-dimensional control of motor units constrained by common inputs spanning one or several muscles.

Low-dimensional neural control of movement has been debated for decades, primarily at the muscle level. In the absence of direct measures of synaptic currents to motor neurons in humans, the correlation (or synchronization) between the firing rates of motor units has been widely used to infer the presence of common inputs (De Luca and Erim, 1994; Keen and Fuglevand, 2004; McIsaac and Fuglevand, 2007; Heckman and Enoka, 2012). In this way, dimensionality reduction methods have been used to estimate the distribution of common inputs to populations of motor units (Madarshahian et al., 2021; Del Vecchio et al., 2023; Levine et al., 2023), with recent evidence that common inputs are distributed to functional rather than anatomical groups of motor units (Hug et al., 2023). In our study, we identified two main latent factors underlying the firing activity of the populations of motor units identified during the reference contractions, regardless of the muscle group (VL-VM or GL-GM). In addition, the angle between motor units (an estimate of common inputs) was not significantly different between GL-GM and VL-VM motor unit pairs, with a notable large variability between motor unit pairs (Figure 2). While this finding aligns with previous work using similar approaches (Del Vecchio et al., 2023), it contrasts with other studies that estimated the degree of common inputs from groups of motor units. Specifically, these studies, which used a coherence analysis have suggested the presence of one dominant common input to the VL and VM muscles (Laine et al., 2015; Avrillon et al., 2021), and a large proportion of independent inputs to the GL and GM muscles (Hug et al., 2021). This discrepancy may be attributed to the level at which each approach infers the degree of common inputs. As coherence analysis is performed on group of motor units, it may not accurately capture the distribution of multiple common inputs to the same motor unit pool (Del Vecchio et al., 2023).

A limitation from previous studies is that observations on the synergistic activity of motor units from one or more muscles have been mostly made from a single motor task. Because the distribution of common inputs could vary with different task constrains, this setup cannot be used to provide definitive proof of hard-wired synergies, and thus of the neural constraints associated with the recruitment of populations of motor units. Here, we leveraged an online motor unit control paradigm, enabling participants to explore different inputs to motor units. Volitional dissociation of GM-GM motor unit pairs was not observed, which aligns with their strong correlated activity during the reference contractions, as indicated by the low angle values (Figure 2). This echoes previous results on a non-compartmentalized muscle (tibialis anterior), in which participants failed to volitionally control individual motor units even after weeks of training (Bracklein et al., 2022). In contrast, most participants were able to dissociate the activity of the GL-GM motor units, which is consistent with their lower correlated activity during the reference contraction (Figure 2). Importantly, this dissociation involved - in some cases (n=4, Figure 3) - volitional recruitment of the higher threshold motor unit without recruiting the lower threshold unit, providing strong evidence of independent control of GL and GM motor units. The fact that no concomitant changes in TA or VL activity were observed during the online feedback tasks (Figure 5) suggests that such independent control was achieved through distinct descending drives rather than focused inhibition. The inability to dissociate the activity of VL-VM motor unit pairs was less expected, especially considering that the dimensionality reduction approach resulted in two latent factors, as observed for GL-GM motor units. Furthermore, the angles between VL and VM motor units were similar to those between GL and GM motor units (Figure 2). Together, these results highlight that the number of latent factors estimated from linear dimensionality reduction methods does not reflect the dimensionality of volitional control of motor units. This is further confirmed by the absence of significant correlation between the angles between motor units used as feedback during the reference contraction and the success rate during the online task (Figure 3). Specifically, some motor units with similar weights in the two latent factors, and presumably a large proportion of common inputs, were dissociated during the online task (e.g. three GL-GM pairs; Figure 3). Conversely, some motor units with different weights in the two latent factors were not independently controlled during the online task.

The inability of the participants to independently control VL and VM motor units despite the relatively large angles between them in the latent space could be explained by the structure of inputs they receive. For example, a single common input could project to the full population of the VL and VM motor units, with heterogeneous projections of inhibitory feedback into this population, leading to decorrelated activity between some units, and thus to the identification of two latent factors. A potential candidate could be recurrent inhibition, whose impact on the decorrelation of activity of neurons receiving common excitatory inputs has been highlighted (Renart et al., 2010; Bernacchia and Wang, 2013). It is important to consider that participants could have achieved GM-GM and VL-VM dissociation if provided with more exposure to the online task, although their success in dissociating the GL-GM motor units suggests that the experimental design was appropriate.

One novelty of this work was to determine whether the dissociation of motor unit activity imposed similar behavior on the motor units that were not displayed in the feedback, and therefore, from which activity was not constrained. Specifically, if non-displayed motor units receive common inputs with those displayed to the participants, they should exhibit similar behavior during online control contractions. This would result in a correlated activity between the displayed and non-displayed units. By extension, we should also observe the same dissociation of activity at the population level as that observed at the paired level. This is what we observed for the non-displayed GL and GM motor units, which exhibited similar behavior to their homonymous motor unit provided as feedback (Figure 4). In addition, except for one pair of motor units (Figure 4), all non-displayed VL and VM motor units remained highly correlated during the online feedback task, mimicking the behavior of the displayed units. Although cortical neural substrates for the volitional selective control of motor units may exist (Marshall et al., 2022), such flexible behavior was not observed when considering GL-GL, GM- GM, and VL-VM pairs of motor units. This is an important finding, providing evidence that despite the dissociation of activity of GL and GM motor units, they were never controlled independently of all others but rather belonged to synergistic groups. In other words, common inputs to motor units from one or several synergistic muscles constrained their activity within low-dimensional manifolds. It is important to note that this result was obtained from a limited subset of motor units located within the same muscle region close to the surface electrodes. Further validation with a more representative sample of motor units, including deep motor units recorded using intramuscular electrodes, is warranted.

Overall, our results suggest that the firing activity of motor units from anatomically defined synergistic muscles is constrained within a low-dimensional control space, with a varying number of dimensions depending on the joint and muscles. The VL and VM motor units seem to receive common inputs, which is consistent with their role in regulating internal joint stress, requiring covariation of their activities (Alessandro et al., 2020). Conversely, because GM and GL muscles have different actions in the frontal plane (Lee and Piazza, 2008), which is supported by our data (Figure 5), the presence of multiple motor common inputs could allow for more flexible control to comply with different task goals, such as joint stabilization. Over time, the central nervous system might compromise between various objectives, such as reducing the control dimensionality (which should involve common inputs to motor neurons) and complying with the task goals in which these muscles are involved (which would require a certain level of flexibility). Future studies should investigate the plasticity of these low-dimensional manifolds when learning new tasks or after nervous system injury.

## Supplemental data availability

The entire dataset (raw and processed data) is available online: https://figshare.com/s/68b5a9b6dfd8e412b01c

## Funding resources

François Hug is supported by the French government, through the UCAJEDI Investments in the Future project managed by the National Research Agency (ANR) with the reference number ANR-15-IDEX-01 and by an ANR grant (ANR-19-CE17-002-01, COMMODE project). Dario Farina is supported by the European Research Council Synergy Grant NaturalBionicS (contract #810346), the EPSRC Transformative Healthcare, NISNEM Technology (EP/T020970), and the BBSRC, “Neural Commands for Fast Movements in the Primate Motor System” (NU-003743).

## Author Contributions

Contribution and design of the experiment: JR, SA, DF, FH; Collection of data: JR; Analysis and interpretation: JR, SA, KT, DF, FH; Drafting the article or revising it for important intellectual content: JR, SA, KT, DF, FH; All authors approved the final version of the manuscript.

